# Daily time constraints limit behavioural capacity to cope with thermally increased metabolic demands

**DOI:** 10.1101/2023.11.06.565854

**Authors:** Evan E. Byrnes, Timo Adam, Carlina C. Feldmann, Larisa Kaplinskaya, Kevin Sticker, Raphael Joshua Fredebeul, Karissa O. Lear, David L. Morgan, Stephen J. Beatty, Roland Langrock, Adrian C. Gleiss

## Abstract

Increased environmental temperatures result in greater energy demands for ectotherms, however, it is currently not clear if these energy demands can effectively be met by increased foraging effort. Here, we tested the temperature dependence of foraging effort and metabolic rate in an aquatic ectotherm across its entire natural thermal range. We developed a novel hidden Markov model to detect behavioural states in long-term body acceleration data collected from free-ranging bull sharks between 19 and 33 °C, and found that increasing temperature altered both the timing and extent of foraging effort. Our data revealed asymmetrical increases of metabolic demands and foraging effort; standard metabolic rates increased exponentially with temperature, but foraging effort increased logarithmically. The observed decoupling of foraging effort and energy demand suggests individuals face increased energy deficits at higher temperatures, confirmed by concomitant reductions in body condition measured in this population with increasing temperatures. We suggest that alterations in the well-established trade-offs between foraging and predation risk coupled with time constraints imposed by high temperatures limit the capacity of animals to cope with environmental temperatures well below critical temperatures.

## Introduction

The rate that animals gain energy is one of the most important factors governing species’ ecology and fitness (Brown et al. 2004). Energy provides the fuel for all physiological processes supporting life (e.g., respiration, nervous function, cellular maintenance), as well as providing the fuel to generate growth, reproduction, and different behaviours (Sibly et al. 2012). The requirement to provide energy to fuel these processes strictly governs animal behaviours, particularly foraging, in that when energy stores are low, animals must forage to maintain body condition and function. While foraging increases energy acquisition, it may also increase risk of predation. As a result, animals adaptively adjust their behaviours to balance these positive and negative effects on fitness (Werner & Anholt 1993; Werner & Hall 1988). However, the rate that animals use energy, and by extension the frequency that animals must forage, is influenced by a myriad of external factors and biological traits that directly impact metabolic rate or indirectly influence the ability of animals to acquire food. Therefore, understanding the influence of environmental factors on species’ ability to forage and meet daily energy demands is critical for species fitness and population persistence under rapidly changing environmental conditions.

In ectotherms, temperature is the predominant environmental factor driving metabolic rate. Typically, standard metabolic rate increases two to three-fold with every 10 °C rise in temperature (Gillooly et al. 2001). As a result, to maintain body condition, ectotherms must increase their energy intake as temperatures rise. When resource limitation is not a constraining factor, animals can meet these increased energy demands by increasing foraging effort. For example, in captive experiments where food is supplied ad *libitum*, foraging rates of fishes and crustaceans increase proportionally to metabolic rate as temperature increases (Baldwin 1957; Cui & Wootton 1988; Elner 1980; Nowicki et al. 2012). However, in the wild, there are several external factors that limit the foraging times and success of animals, and as a result, may curtail their ability to meet their increased energy demands at higher temperatures.

Obtaining enough food to meet energy needs while simultaneously trying not to be eaten is a pervasive trade-off that limits the timing of foraging for many organisms, including lower-level predators (i.e., life-dinner principle, Dawkins & Krebs 1979). Animals tend to limit foraging to times when predators are less abundant or when the likelihood of predation is reduced, such as during the day when predators may be more easily detected visually (Heithaus & Dill 2006; Lima & Dill 1990). However, animals may prefer to forage during a specific time of the day when foraging efficiency can be maximized due to species-specific traits that are optimized under specific conditions (Bosiger & McCormick 2014; Erkert 2000; Hammerschlag et al. 2017). For example, animals that display nocturnal foraging patterns possess adaptations that enhance sensory capabilities in low-light conditions, such as retroreflective optical tissues (Gruber 1977), electrical sensory abilities (Kramer 1996), or pressure receptive cells (Bleckmann & Zelick 2009). Combined, variation in predation risk and a species’ ability to capture prey during certain periods in the day, among other factors, likely constrain the capacity of free-ranging animals to increase the amount of time they spend foraging or capacity to shift their temporal foraging niches.

This study aimed to test how temperature induced increases in metabolic rate affect the duration and timing of foraging behaviour in an ectotherm experiencing drastic seasonal temperature changes. We applied hidden Markov models to biotelemetry-derived activity data from an ectothermic mesopredator (juvenile bull sharks, *Carcharhinus leucas*) within the Fitzroy River, Kimberley, Western Australia to infer temporal variation in foraging effort (i.e., time spent foraging). The Fitzroy River serves as a “natural mesocosm” to explore the effects of temperature on species physiology and ecology because extreme fluctuations in river flow cause animals to be seasonally constrained to small pools, where they are forced to endure increases in water temperatures (Whitty et al. 2017). As water temperatures increased, we hypothesized that the overall amount of time individuals spent foraging each day would increase, with temporal foraging niches expanding overnight when temperatures, and therefore metabolic costs of activity are lowest.

## Methods

### Study site

This study was conducted within the freshwater reaches of the Fitzroy River, located in the Kimberley region of Western Australia, Australia. This environment is highly dynamic, experiencing environmental extremes with rainfall restricted to a distinct wet season (approximately mid-December to May). During the wet season, monsoonal rains fill the river catchment and the river is fully connected; during the dry season, flow drops and in most years eventually ceases, forming separated pools that are fed only by local aquifers (CSIRO 2018). Within these pools aquatic animals become trapped until flow increases again in the following wet season and are therefore forced to cope with prevailing environmental conditions, making this an ideal natural system for studying the ecophysiology of wild animals. Throughout the dry season, mean daily water temperature gradually increases from around 18 °C to approximately 35 °C (Lear et al. 2020; Whitty et al. 2017). This study was conducted in two different pools over 100 km upstream from the mouth of the river: Myroodah Pool and Camballin Pool (see Whitty et al. 2017).

### Capture and tagging

Bull sharks (Carcharhinus leucas) were captured from July to August in both 2015 and 2016 from a 4-m motorized skiff, using gillnets (152-202 mm stretched mesh). Once captured, sharks were held horizontally next to the skiff, and the sex of each animal was recorded and the total length (TL) and girth were measured. Two acoustic transmitters (V13T & V13AP; Vemco, Innovasea, NS, CAN) were surgically implanted into the peritoneal cavity of sharks through a small incision anterior to the pelvic fins, and sharks were released. V13T tags transmitted a temperature reading (°C; accuracy: ±0.5 °C, resolution: 0.15 °C) and V13AP tags alternately transmitted body acceleration data (m/s^2^) and depth data (m; accuracy: ±1.7 m, resolution: 0.08 m). For each body acceleration transmission, triaxial acceleration was recorded at 10 Hz for a 20-s period and was processed within tags and transmitted as mean vectorial dynamic body acceleration (VeDBA). All tags transmitted with a random nominal delay between 120 to 180 s, to reduce overlap of transmissions between tags. Transmissions were recorded by acoustic receivers (VR2W; Vemco, Innovasea, NS, CAN) placed approximately 500 m apart throughout the length of each pool. Receivers had a detection range of 300-400 m (Whitty et al. 2017). All animal use was conducted under a Murdoch University Animal Ethics permit (no. RW2757/15).

### Data pre-processing

All data processing and analyses were conducted in R (version 3.5.2., R Core Team 2019). Prior to analysis, false detections were removed from the dataset. Detections of a given ID were considered false if they were recorded only once on a given receiver within a one-hour period or when subsequent detections recorded by different receivers were too close in time for an individual to reasonably travel the distance separating the receivers. Additionally, some sharks appeared to have died due to natural causes (i.e., predation or starvation) or expelled tags before the end of tag battery life, indicated by consistent depth transmission and zero body acceleration. Therefore, data after apparent death or tag expulsion were excluded prior to analysis. Detections were then pooled into hourly bins, and the mean hourly VeDBA, temperature, and depth for each individual were calculated for the duration of deployments.

### Behavioural pattern

To examine how foraging patterns changed as a function of increasing water temperature, acceleration data were analysed using a hidden Markov model (HMM). HMMs are a class of stochastic time-series models that allows an observed time-series (e.g., acceleration time-series) to be decoded into the most likely sequence of underlying, hidden states. These states can be considered as noisy measurements of behaviours, which cannot be explicitly observed (Leos-Barajas et al. 2016). Predicted state sequences can then be decomposed to compute the probability of animals being in different states as a function of various covariates. HMMs have widely been used in ecological research to decode different movement modes (e.g., transient vs resident) from biotelemetry recorded time series of spatial location data, however, they are more generally applicable for classifying observations from various types of time-series data into different behavioural states, including mark-recapture, consecutive diving behaviours, and biologger data (e.g., acceleration, Zucchini et al. 2016). Here, we used HMMs to decode hourly time series of acceleration data (i.e., VeDBA) from acoustic tags into relative activity states, as proxies for different behaviours. We additionally modelled the probability of sharks switching between activity states as a function of time of day and average daily water temperature.

We developed a two-state HMM, where sharks could be in either a low state of activity (state 1) or a high state of activity (state 2), based on mean hourly VeDBA values from acoustic tags. Hourly VeDBA observations were assumed to be conditionally independent given the states, using Gamma state-dependent distributions. The state process was assumed to be governed by a first-order, N- state Markov process (i.e., current state is temporally correlated only with the previous state). The evolution of the activity states over time was governed by a non-homogenous transition probability matrix, Γ, defined as:

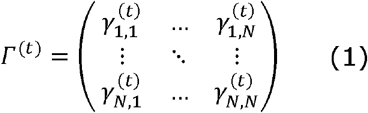

where 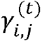 is the probability of transitioning from state i to state j at time t. Using row-wise multinomial logit links, the entries of the transition probability matrix were calculated as a function of two covariates: average daily temperature and hour of the day. Temperature was averaged over each day before inclusion as a covariate because the variation in hourly temperature intrinsically comprises seasonal and daily variation. This reduction of temperature into a daily timescale was important for two reasons: 1) it reduced the information of the temperature covariate to seasonal variation; and 2) avoided problems with multicollinearity between time of the day and temperature, as the daily variation in temperature was captured by the temporal covariate, hour of the day. Hour of the day was included using four trigonometric functions: two corresponding to a 24-hour cycle (sin(*2πt/24*), cos(*2πt/24*)) and two corresponding to a 12-hour cycle (sin(*2πt/12*), cos(*2πt/12*)). As we were primarily interested in how diel behavioural patterns evolved as a function of seasonal temperature, interactions between daily temperature and all four hour-of-the-day trigonometric functions were also included. Accordingly, entries of the transition probability matrix were a function of the following equation:

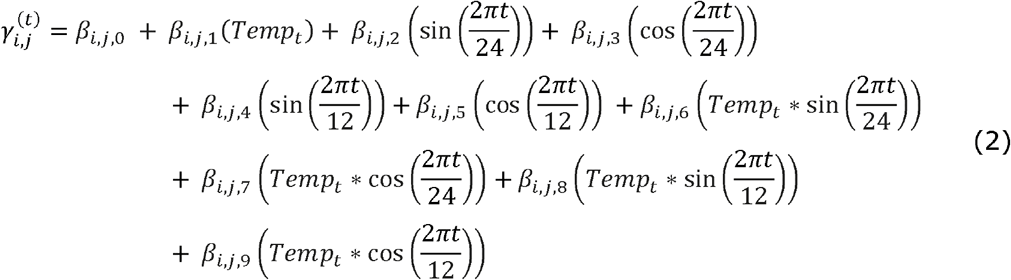

with *β*_*i,j,k*_ denoting the coefficient of the k-th additive term.

To account for differences in VeDBA between individuals that may occur as a result of different body sizes (Gleiss et al. 2011; Qasem et al. 2012), separate parameters for the state-dependent Gamma distributions (i.e., mean and variance) were estimated for each individual. Additionally, temperature has a pronounced influence on the physiology and biomechanical performance of ectotherms, typically described by a bell-shaped thermal performance curve (Angilletta Jr et al. 2002; Rall & Woledge 1990). Consequently, the rate of body movement associated with behaviours can intrinsically increase with temperature, causing higher acceleration values, without the animal necessarily changing the frequency of occurrence of different behaviours. Therefore, to prevent from overestimating the amount of time spent within a behavioural state, it is crucial to account for such thermally associated changes in kinematic performance. To accommodate these temperature-varying state accelerations within our model, individual-specific (and state-dependent) VeDBA means were expressed as nonparametric functions of time of the year modelled using P-splines (using 15 B-spline basis functions of 4^th^ order, and smoothing parameters chosen subjectively based on visual inspection of the goodness of fit). The model thus is a special case of a Markov-switching generalised additive model (Langrock *et al*. 2017). The same transition probability matrix was used across all individuals, assuming the same effect of covariates on all individuals.

HMMs were fit using numerical maximum likelihood estimation using the nlm optimizer in R (R Core Team 2019), with 1000 iterations run using randomized starting parameter values to ensure global maxima were identified. Once the final model was established, the Viterbi algorithm was applied to decode the most likely state sequence underlying each individual VeDBA time series, given the estimated individual state-dependent distributions and covariate dependent state transition probabilities (Zucchini *et al*. 2016). Lastly, to inspect the probability of sharks being in each activity state as a function of hour of the day and temperature, the marginal probabilities of states were determined from the stationary distributions at all observed combinations of covariate values (Patterson *et al*. 2009).

To determine how sharks increased foraging effort as a function of temperature, we examined the relationship between the time spent foraging each day and the daily water temperature. Daily percentage time foraging was calculated as a proxy of daily foraging effort based on Viterbi sequences estimated for each individual from HMMs. Percentage time foraging was calculated as the number of hours that an individual was estimated to be in the high-activity (i.e., foraging) state each day divided by 24 hours. Daily estimates of percentage time foraging for each individual were grouped into one-degree temperature bins, based on mean daily temperature. The mean percentage time spent foraging was estimated for each individual within each bin. The relationship between daily percentage time foraging and temperature was then explored using a logistic growth curve.

To determine how foraging effort changed with metabolic demand, daily energy expenditure was estimated for each individual using the predictive equation established by Lear et al. (2020):

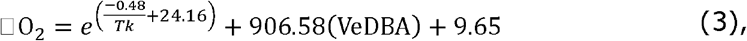

where *T* is the body temperature in kelvin and *k* is Boltzmann’s constant (8.62×10^−5^ eV K^−1^).

Mean hourly temperature and mean hourly VeDBA recorded by tags for each individual were input into the equation to estimate mean hourly energy expenditure. For each day, the mean hourly energy expenditure of individual was estimated and multiplied by 24 hours to represent mean daily energy expenditure. To allow comparison between individuals, although most individuals did not substantially vary in size (Table 1), mass-specific daily energy expenditure estimates (mg O_2_ kg^-0.86^ day^-1^) were used.

**Table 1.**
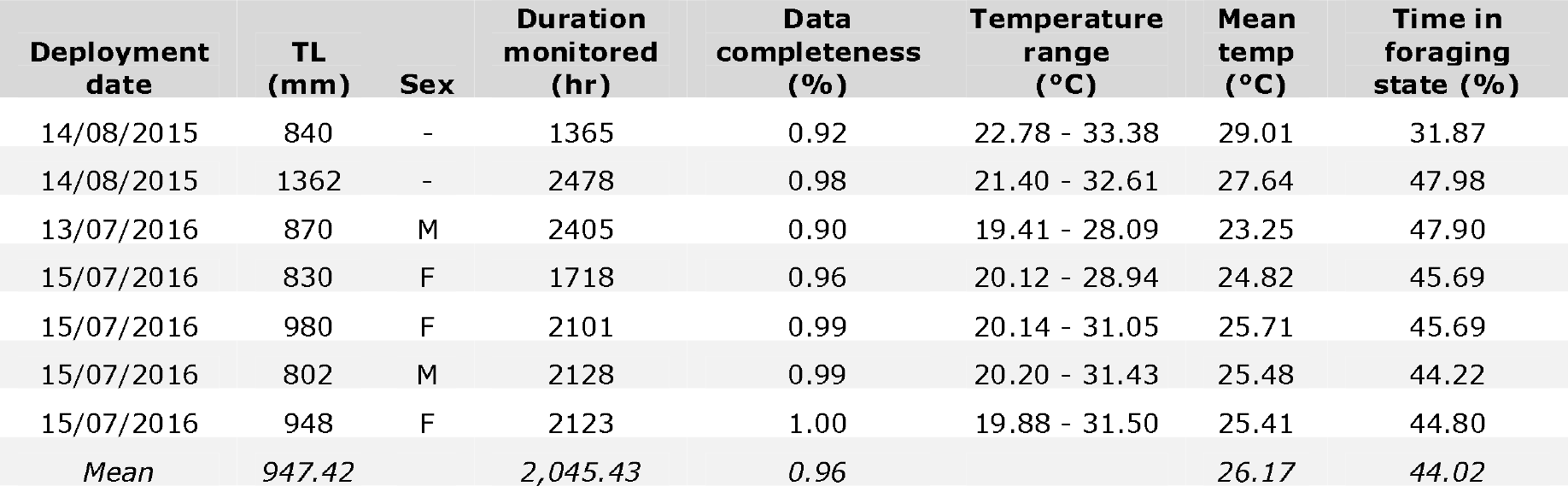
Deployment summary for sharks used in this study.

| Deployment date | TL (mm) | Sex | Duration monitored (hr) | Data completeness (%) | Temperature range (°C) | Mean temp (°C) | Time in foraging state (%) |
| --- | --- | --- | --- | --- | --- | --- | --- |
| 14/08/2015 | 840 | - | 1365 | 0.92 | 22.78 - 33.38 | 29.01 | 31.87 |
| 14/08/2015 | 1362 | - | 2478 | 0.98 | 21.40 - 32.61 | 27.64 | 47.98 |
| 13/07/2016 | 870 | M | 2405 | 0.90 | 19.41 - 28.09 | 23.25 | 47.90 |
| 15/07/2016 | 830 | F | 1718 | 0.96 | 20.12 - 28.94 | 24.82 | 45.69 |
| 15/07/2016 | 980 | F | 2101 | 0.99 | 20.14 - 31.05 | 25.71 | 45.69 |
| 15/07/2016 | 802 | M | 2128 | 0.99 | 20.20 - 31.43 | 25.48 | 44.22 |
| 15/07/2016 | 948 | F | 2123 | 1.00 | 19.88 - 31.50 | 25.41 | 44.80 |
| *Mean* | *947.42* |  | *2,045.43* | *0.96* |  | *26.17* | *44.02* |

### Vertical movement

To examine how vertical movement of sharks was influenced by temperature throughout the year, we used generalized additive mixed models (GAMMs, “mgcv” package, Wood & Wood 2015). Average hourly depth was modelled as a function of hour of the day and temperature, with individual ID included as a random effect. Hour of the day was fit using a cyclic cubic regression spline, whereas temperature was included in models via an interaction with time of day because we were primarily interested in how the diel vertical movements pattern of sharks changed as a function of increasing daily temperature. An autoregressive correlation structure at time lag=1, nested within individual, was also included in models to account for serial autocorrelation in depth. Models with all possible combinations of covariates were constructed, and compared based on corrected Akaike’s Information Criterion (AICc), with a decrease ≥2 considered as an improvement in model fit (Zuur et al. 2009). All models were constructed using a Gaussian error distribution, and the final model fit was assessed based on inspection of residual plots.

## RESULTS

### Deployments

Between 2015 and 2016, acoustic transmitters were deployed in 16 bull sharks, however, data from only seven sharks was sufficient for analyses. Imputation of more than 10% of data in time series data may introduce substantial bias into analyses, such as HMMs (Zucchini et al. 2016). Therefore, all animals with less than a <90% detection rate were excluded from analysis, leaving seven individuals (Table 1). These seven individuals were monitored for a total of 14,318 hours (mean: 2,045.43±387.84 hrs), or over 602 days (mean: 86±16.44 days) (Table 1). Mean hourly temperature ranged from 19.41 °C to 32.73 °C (mean: 26.13±3.37 °C), with mean temperature increasing throughout the duration of the deployments (Fig. 1). Hourly mean VeDBA values ranged from 0.10 m/s^2^ to 4.24 m/s^2^ (mean: 0.71±0.25 m/s^2^) and generally increased throughout the dry season (Fig. 1). Hourly mean depth use varied from 0.00 m to 4.01 m (mean: 0.79±0.61 m, note: tag mounted on dorsal fin allowing tag to record depth of 0 m while majority of shark remains submerged).

**Figure 1.**
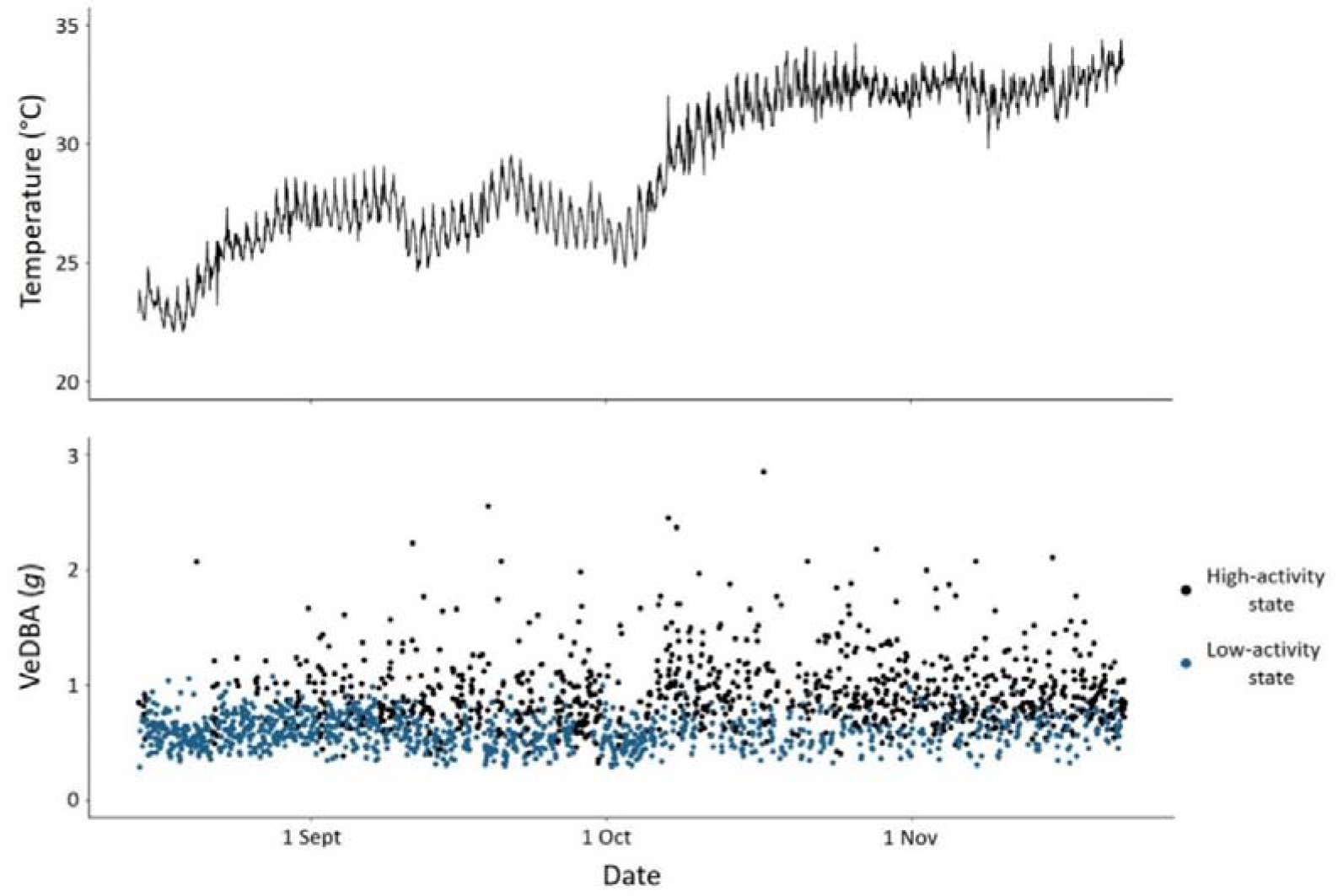
Time-series of the mean hourly temperature and VeDBA for the whole deployment from a single shark. VeDBA is coloured by activity state, which was decoded via the Viterbi algorithm.

### Behavioural pattern

The final hidden Markov model had two states, which were interpreted as different activity states associated with different behavioural modes. The first state had a lower mean and standard deviation of log(VeDBA), hereafter called the “resting state”, and was inferred to represent resting or slow swimming associated with routine behaviours. The second state had a higher mean and greater standard deviation of log(VeDBA), hereafter called the “foraging state”, and was inferred to represent fast and burst swimming associated with prey searching and foraging behaviours. These interpretations are similar to those of other studies that have identified activity states of fish based on acceleration data (Byrnes *et al*. 2021; Papastamatiou *et al*. 2018).

Sharks demonstrated a clear diel cycle in the activity-state pattern, with the probability of the foraging state generally lowest during the day, and highest during the night or early morning. However, the timing and duration of periods when sharks were most likely to be in the foraging state shifted as temperature increased (Fig. 2). When daily temperatures were lower, sharks demonstrated a nocturnal activity pattern, where the probability of the foraging state peaked overnight and was lowest during the day. At higher temperatures, sharks demonstrated more of a crepuscular activity pattern. Notably, as temperature increased, the duration of the resting period during the day decreased and simultaneously shifted to later in the afternoon. In contrast, as temperature increased, the duration of the foraging period increased and shifted to later in the night, and eventually included morning. For instance, at 20 °C, the probability of the foraging state peaked for about an hour from 02:00 to 03:00 and was lowest for multiple hours from 08:00 to 16:00. At 31 °C, the probability of the foraging state was highest for multiple hours from 04:00 to 09:00 and was lowest for approximately an hour from 16:00 to 17:00 (Fig. 2).

**Figure 2.**
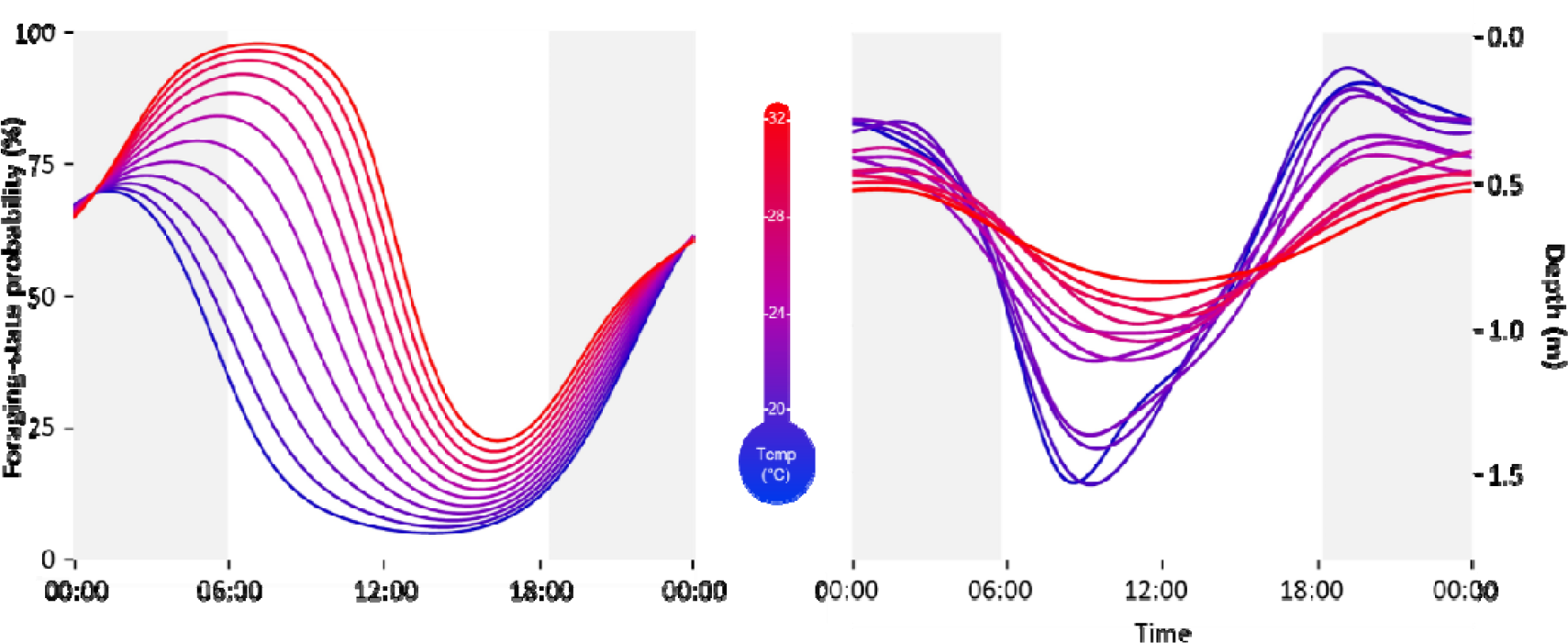
Daily patterns of the probability of being in the high-activity state (i.e., foraging state; left) and mean hourly depth (right) as a function of daily temperatures. Lines are coloured by temperature from 20 to 31 °C, represented by the transition from blue to red. Daily light periods are indicated by background shading on plots, with the grey areas indicating nocturnal periods, which were determined based on the average sunset and sunrise times over the entire study period.

Overall, sharks spent a mean of 44.02% of the time in the foraging state, with little individual variation, apart from one individual (Table 1). However, sharks spent relatively more time in the foraging state as daily water temperature increased through the season (Fig. 1, 2), shifting from the majority of time spent in the resting state at cooler temperatures to the majority of the time spent in the foraging state at warmer temperatures. Sharks spent an average of 12.69% of the day in the foraging state at 20 °C, increasing to 63.44% of the day at 31 °C (Fig. 3). Notably, sharks demonstrated a nonlinear increase in the amount of time spent foraging each day (Table A1; Fig. 3), where foraging increased logarithmically, eventually plateauing at temperatures above approximately 27 °C. This contrasted with field metabolic rate, which increased exponentially with temperature.

**Figure 3.**
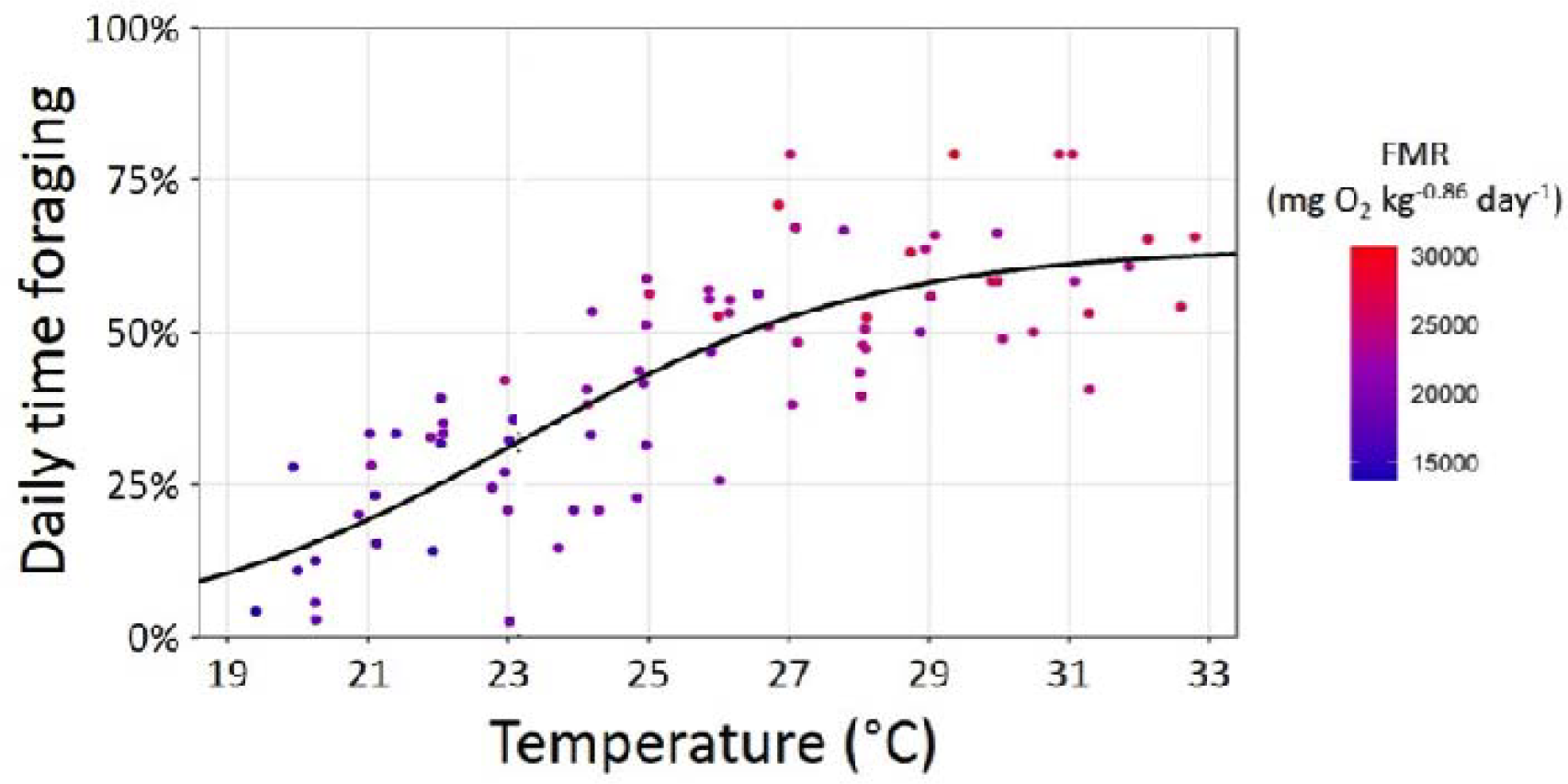
Mean daily proportion of time spent foraging each day for each individual as a function of temperature. Estimates of the percentage of time spent foraging each day were grouped into one-degree temperature bins based on mean daily temperature, and the mean for each bin for each individual was estimated. Each point represents an individual within a temperature bin and is coloured by mean daily FMR observed for an individual within a temperature bin.

### Vertical movement

The final GAMM describing daily depth use patterns included an interaction between hour of the day and average daily temperature, with individual ID as a random effect (Table 2). Sharks demonstrated clear diel vertical movements throughout the season, using deeper waters during the middle of the day and shallower waters during the nocturnal period (Fig. 2). However, the diel pattern was more pronounced at lower water temperatures, with substantial differences observed between depth use in the day vs night during cooler periods, which decreased as daily water temperature increased. Specifically, early in the dry season, at daily mean temperatures of 19 °C, mean hourly depth varied by nearly 1.5 m throughout the day. In comparison, at temperatures near 31 °C, mean hourly depth varied by less than 0.5 m throughout the entire diel period (Fig. 2). It is important to note that during most years, the maximum depth throughout most of the stretch of river pools is around 2 m, with an absolute maximum depth of about 5 m. The timing of shallowest and deepest depths occupied also changed as daily water temperature increased (Fig. 2). As temperature increased, the period of shallowest depth use gradually shifted from evening (≄21:00 at 20 °C) to early morning (≄05:00 at 32 °C), whereas the shift in deepest depth use was relatively smaller, from late-morning (≄10:00 at 20 °C) to mid-afternoon (≄15:00 at 32 °C).

**Table 2.**
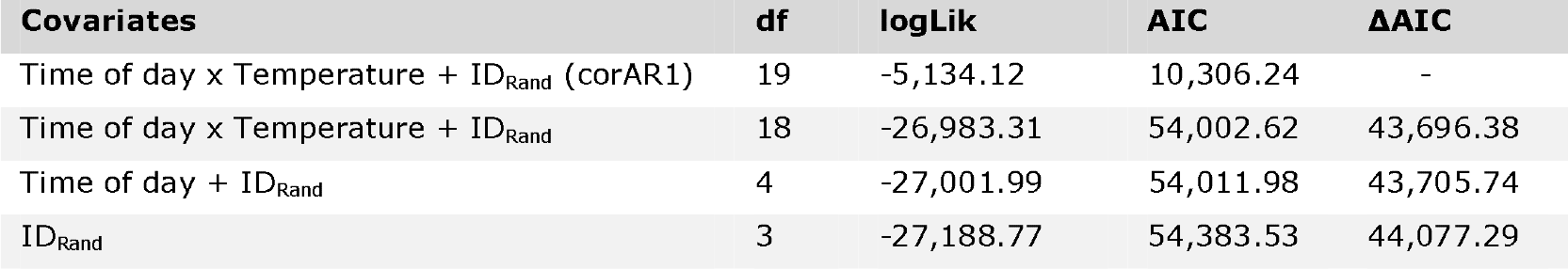
Depth use GAMM selection table. Models ranked by AIC. ‘x’ indicates an interaction between terms.

| Covariates | df | logLik | AIC | ΔAIC |
| --- | --- | --- | --- | --- |
| Time of day x Temperature + ID<sub>Rand</sub> (corAR1) | 19 | -5,134.12 | 10,306.24 | - |
| Time of day x Temperature + ID<sub>Rand</sub> | 18 | -26,983.31 | 54,002.62 | 43,696.38 |
| Time of day + ID<sub>Rand</sub> | 4 | -27,001.99 | 54,011.98 | 43,705.74 |
| ID<sub>Rand</sub> | 3 | -27,188.77 | 54,383.53 | 44,077.29 |

## DISCUSSION

For the first time, this study showed that increases in the foraging effort of a free-ranging ectotherm do not necessarily increase proportional to metabolic demands. To grow and survive in changing environments, animals should increase foraging to meet temperature induced exponential increases in their metabolic demands, and as such foraging effort should also increase exponentially with temperature. However, our results demonstrated that as temperature increased, the amount of time invested in foraging increased asymptotically, causing a temperature induced decoupling of energy intake and output.

This decoupling has the ramification of changing net energy intake as a function of temperature. In ectotherms, both metabolic rate and maximum energy intake are known to be temperature dependent, however, their responses to temperature are markedly different, causing net energy gain to be temperature dependent. Englund et al. (2011) have shown the temperature dependency of energy intake to be steeper at lower temperatures than metabolic rate, but the difference is stabilized toward intermediate temperatures, and at higher temperatures, the temperature dependency of energy intake exhibits a shallower and opposing response to that of metabolism. The resultant effect of this pattern is a bell-shaped thermal growth response (Brett et al. 1969). Given that foraging time should generally be proportional to energy gain (Calcagno et al. 2014), it is expected that thermal dependence of foraging time should follow the intake pattern predicted by Englund et al. (2011; but see Grigaltchik et al. 2012).

In this system, bull shark foraging effort appeared to follow a logarithmic response to temperature, with the rate of increase in effort gradually slowing to a plateau between 27 to 30 °C. In contrast, energy expenditure is known to continually increase exponentially in bull sharks at temperatures well above this range (Lear *et al*. 2019). Assuming that energy intake is proportional to the time invested in foraging, the proportionally slower increase in energy intake than metabolism at high temperatures likely causes animals to face energy deficits at higher temperatures, which if prolonged could lead to starvation. Indeed, the same sharks within this system have been shown to experience approximately a 30% loss in body mass between July and December (Lear et al. 2020). Nevertheless, we could not determine if the energy loss may have also been influenced by changes in prey abundances throughout the season. To do so would require coupling estimates of daily energy expenditure with estimates of daily consumption rates provided by continual measurements of intake or frequent measurements of body condition combined with bioenergetics modelling and regular prey species abundance estimates

In addition to meeting energy demands, animals have several daily requirements, for instance, having a threshold for the amount of time needed for rest (Zeppelin & Rechtschaffen 1974) and the need to shelter to avoid direct predation (Lima & Dill 1990) or physiologically challenging environmental conditions (e.g., high afternoon temperatures, Sunday et al. 2014). However, the trade-offs governing how animals partition their time to meet these requirements may change with temperature. For example, deciding when and how long to forage is commonly influenced by a trade-off between the risk of starving versus the risk of predation (Werner & Anholt 1993). At low temperatures, when metabolic rates are low, animals should be less willing to forage while faced by predation risk, whereas at high temperatures there will be a relative increase in individuals’ willingness to forage despite this risk, and as such increases in the time spent foraging (Abram et al. 2017; Lima & Dill 1990).

Time foraging, however, cannot increase boundlessly, as certain periods of the day may be unusable, owing to physiological limits. As water temperature increased throughout the year, animals were likely faced with potentially damaging temperatures that limited the duration and timing of foraging behaviours, particularly in shallow foraging areas where temperatures can reach above 34 °C (CSIRO 2018; Lear et al. 2020). While the upper critical temperature for bull sharks has not yet been identified, similar temperatures have been reported to be lethal in juveniles of other shallow water tropical elasmobranch species (Bouyoucos et al. 2021). Furthermore, these temperatures are above the optimum for locomotory performance and therefore the ability to capture prey may be diminished when foraging in waters at or above this temperature (Grigaltchik et al. 2012; Lear et al. 2019). Even if temperatures do not reach lethal levels, the energetic cost of highly active behaviours at such high temperatures likely outweigh possible energy gains, making foraging during the hottest periods of the day energetically unjustifiable. For instance, nocturnal activity of animals in tropical and desert environments are often associated with such physiologically imposed thermal limits (Kronfeld-Schor & Dayan 2003; Murray & Smith 2012). Alternatively, but not exclusively, as the amount of time invested in active behaviours such as foraging increases, it is possible that animals may approach a limit imposed by minimum daily resting requirements (Zeppelin & Rechtschaffen 1974). Therefore, the asymptotic relationship observed here could be a product of thermally induced shifts in the trade-offs mediating behavioural decisions, with risk associated constraints at lower temperatures followed by critical physiological limits at higher temperatures.

The asymptotic temperature dependency of the time devoted to foraging demonstrated here is likely caused by constraints imposed by ecological mechanisms that are common to virtually all taxa. Of course, the exact properties of this relationship will intrinsically vary across species, owing to differences in the relative level of predation risk and physiological traits. For example, the relative increase in the amount of time foraging each day with increased temperature may be more gradual in species that experience lower levels of predation because foraging at lower temperatures may only be constrained by limits of digestive rates. Additionally, the amount of time foraging may plateau more gradually or at higher temperatures in species with relatively lower metabolic rates because they may more easily meet metabolic demands in shorter periods of foraging, and therefore may not be time constrained until higher temperatures. This would explain why freshwater sawfish (*Pristis pristis*) within this system, which have substantially lower standard metabolic rates and temperature sensitivity than bull sharks, were found to lose relatively lower percentages of body mass over the dry season (Lear *et al*. 2020). In addition, foraging strategies and whether or not environmental temperatures within foraging areas reach damaging levels could further influence how time investments into foraging change in response to elevated daily temperatures.

Our unique development of a hidden Markov model allowed the classification of long-term acoustic telemetry derived activity data, where periods when sharks were in a high-activity state were interpreted as foraging periods. It is possible that occurrence of exceptionally high-activity behaviours, such as predator evasion responses could cause overestimation of the time spent in a foraging state. However, such behaviours are generally short-lived (≤1 minute), and thus were likely to substantially influence acceleration values averaged over long durations (hourly). However, an inherent problem remains with using unsupervised classification algorithms such as HMMs, in that it can be difficult to determine the behaviours associated with each identified state without validation of data from direct observations (Leos-Barajas *et al*. 2016; Patterson et al. 2019). Despite this, it is known that carcharhinid sharks regularly swim at higher swim speeds, and thus increase associated acceleration values while searching and foraging (Papastamatiou *et al*. 2015; Roemer *et al*. 2016; Watanabe *et al*. 2019). In addition, times spent in shallow environments where prey species were commonly observed (e.g., mullet and other small fishes) was synonymous with increased times in the high activity state. Although foraging may occasionally occur at depth, sharks in this system likely forage over the shallow flats, where trapping prey against the shoreline aids in their capture (personal observation). Therefore, we are confident that the high activity state identified by our HMM represented foraging associated behaviours, including searching and capturing of prey.

Comparative physiological research has provided a thorough understanding of the critical limits to species’ function and performance, leading to major concern about the survival of wild populations and ecosystem stability as changes in environmental conditions approach these limits. For example, under current climate projections, environmental temperatures are expected to increase above the physiological optimal temperatures of many tropical ectotherms by 2100, leading to widespread extirpations (Nguyen et al. 2011; Tewksbury *et al*. 2008). However, the results of this study indicated that ecological constraints on behaviour can limit the fitness and survival of free-ranging animals well before such experimentally identified critical environmental limits are reached. Here, we clearly demonstrate that predation risk and time constraints can impose powerful limits on the foraging of an ectotherm, hindering their ability to meet energy requirements as temperatures increase. These results highlight that behavioural plasticity is likely a fundamentally limiting aspect of species capacity to cope with environmental change, especially in oligotrophic systems and/or in species with a high-energy demand. As such, advancing our understanding of the constraints on the behavioural plasticity of free-ranging animals is critical to comprehensively understanding how species respond to environment change and for accurately predicting their survival and population persistence under challenging environmental conditions.

## Acknowledgements

We would like to thank the Nyikina Mangala Rangers, J. Whitty, and B. Bentley for help with data collection in the field. We thank the Nyikina Mangala people, the original custodians of the land on which this work was conducted, and we pay our respects to their Elders, past and present. Fieldwork was generously funded by grants from the Australian Research Council Discovery Early Career Research Award (Project number 150100321), the Fisheries Society of the British Isles, Australia Pacific Science Foundation, the Waitt Foundation, Western Australian Government State Natural Resource Management Program and Murdoch University. Animal use was conducted under a Murdoch University Animal Ethics permit (RW2757/15).

---

**Table A1.** Model selection table for models examining influence of temperature on foraging effort (i.e., daily time spent foraging). For simplicity, model formulas not shown; all models were fit with temperature as the explanatory variable.

| model | df | AICc | log-likelihood |
| --- | --- | --- | --- |
| Linear | 78 | -107.49 | 56.90 |
| 2<sup>nd</sup> order polynomial | 77 | -113.73 | 61.13 |
| Nonlinear least squares | 77 | -114.30 | 61.41 |

